# xTrimoGene: An Efficient and Scalable Representation Learner for Single-Cell RNA-Seq Data

**DOI:** 10.1101/2023.03.24.534055

**Authors:** Jing Gong, Minsheng Hao, Xin Zeng, Chiming Liu, Jianzhu Ma, Xingyi Cheng, Taifeng Wang, Xuegong Zhang, Le Song

## Abstract

The advances in high-throughput sequencing technology have led to significant progress in measuring gene expressions in single-cell level. The amount of publicly available single-cell RNA-seq (scRNA-seq) data is already surpassing 50M records for human with each record measuring 20,000 genes. This highlights the need for unsupervised representation learning to fully ingest these data, yet classical transformer architectures are prohibitive to train on such data in terms of both computation and memory. To address this challenge, we propose a novel asymmetric encoder-decoder transformer for scRNA-seq data, called xTrimoGene, which leverages the sparse characteristic of the data to scale up the pre-training. This scalable design of xTrimoGene reduces FLOPs by one to two orders of magnitude compared to classical transformers while maintaining high accuracy, enabling us to train the largest transformer models over the largest scRNA-seq dataset today. Our experiments also show that the performance of xTrimoGene improves as we increase the model sizes, and it also leads to SOTA performance over various downstream tasks, such as cell classification, perturb-seq effect prediction, and drug combination prediction.

## 1 Introduction

Recently, Artificial Intelligence (AI) technology has demonstrated promising results for addressing scientific problems. This AI4Science (AI for Science) paradigm witnessed diverse successful biological and pharmaceutical applications, including protein analysis (Jumper et al., 2021; Lin et al., 2022; Zhang et al., 2022; Xiao et al., 2021; Brandes et al., 2022), RNA modeling (Chen et al., 2022a), and genomics modulation (Pesaranghader et al., 2022). However, most existing AI models have predominantly focused on protein sequences, neglecting the growing volume of high-throughput experimental sequencing data in the form of gene expression values. Single-cell RNA sequencing (scRNA-seq) technology has transformed the field of cell biology and enabled us to understand cell-cell, cell-gene and gene-gene relations at individual cell level (Jovic et al., 2022; Chen et al., 2019). This technique determines the expression levels of thousands of genes in parallel, and has been shown useful in the study of cellular heterogeneity and the identification of unique molecular signatures within a large population (Chen et al., 2022b; Li et al., 2022). This unveiled information is crucial for understanding complex biological systems and disease progression (Jovic et al., 2022; Chen et al., 2019). Integrating and modeling such large-scale scRNA-seq data can reveal rich cellular information and benefit various biological task learning.

Representation learning from scRNA-seq data (Flores et al., 2022) has been an active area of research in past decades. For example, scVAE (Grønbech et al., 2020) and scVI (Lopez et al., 2018) apply a variational autoencoder framework to derive low-dimensional cell embeddings, cscGAN (Mohamed Marouf, 2020) uses a Generative Adversarial Network (GAN) architecture to generate cell-type specific expression profiles, and SAVER-X (Wang et al., 2019) is capable of removing batch effects across datasets. Despite the success of these customized algorithms, they tend to be computationally inefficient and labor-intensive. This prompts us to explore a general-purpose model that first learns underlying knowledge from scRNA-seq data and then generalize it to different tasks in a unified manner. We draw inspiration from the pre-training and fine-tuning paradigm in Natural Language Processing (NLP), which has shown great success in improving various downstream NLP task performance (Qiu et al., 2020; Han et al., 2021; Hendrycks et al., 2019). The efficacy of the pre-training technology is highly correlated with model size and data scale, following a power-law relationship (Kaplan et al., 2020). In light of these findings, we aim to investigate the potential of applying similar approaches to representation learning in scRNA-seq data. The first published pre-trained model for single-cell data is scBERT, which uses a low-rank transformer (Yang et al., 2022) to analyze the scRNA data. It learns the cellular representation by randomly masking a percent of non-zero gene expression values and tries to recover them. scBERT has achieved state-of-the-art results for cell-type annotation tasks. The study shows the potential of pre-training strategy for single-cell biology research.

However, scBERT has certain limitations in fully utilizing scRNA-seq data properties. These limitations include: (1) Scalability. The core of the scBERT algorithm involves the incorporation of a series of attention blocks to capture gene-gene interactions. In spite of employing a faster Performer architecture, the computational complexity remains substantial due to the large number of genes (almost 20,000) and the sparsity of scRNA-seq data, with nearly 90% of values being zero, leading to many redundant computations (e.g., self-attentions between zero tokens). These characteristics result in a high computational cost, requiring approximately 2.65 × 10^19^ FLOPs to train 5 million samples over 5 epochs, which equals almost 20 days of training on an A100 GPU for only a 8.9 million parameter scBERT model. (2) Limited resolution for expression values. In order to make use of the Performer architecture, the gene expression values are discretized into integer values via a rounding operation. The binning process limits the model’s ability to distinguish closeness and similarity between gene expression values. For example, two close values are potentially mapped to separate embeddings (e.g., 1.99 and 2.01 are mapped to 1 and 2), and two distant values are potentially mapped to identical embeddings (e.g., 1.99 and 1.01 are mapped to 1 and 1). The strategy leads to a loss of resolution and introduces bias during model training, resulting in sub-optimal performance.

In order to address the challenges associated with scRNA-seq data modeling and consider the unique nature of this data (as discussed in Section 2), we present a novel and efficient framework, xTrimoGene, for pre-training large-scale scRNA-seq data. Our framework makes the following key contributions:

- We design an asymmetrical encoder-decoder architecture to guide the pre-training process, which enables us to learn a high-capacity model for single-cell RNA-seq data. Our model achieves an improvement in the speed of pre-training of over 3 times compared to previous decoder-only models such as scBERT.
- We illustrate that the efficiency and scalability of our model allow us to train the largest single-cell pretrained model to date, with approximately 100 million parameters for the xTrimoGene-100M model, using a scRNA-seq dataset of approximately 50 billion effective gene tokens that we curated from public data sources.
- The pre-trained model xTrimoGene achieved remarkable results in multiple downstream tasks, including cell type annotation, perturbation response prediction and synergistic drug combination prediction.

## 2 Characteristics of Single-Cell RNA-seq Data

scRNA-seq generates a large, sparse expression matrix, where each row represents a cell (sample) and each column a gene (feature). This dataset presents several challenges, and requires a specialized architecture to effectively model the data.

First, approximately 20,000 genes (columns) are shared across cells. In contrast to the corpus in NLP, the genes can be arbitrarily reordered. The relation between genes depends on biological pathways rather than local contexts, where the latter shapes spatial information in Computer Vision (CV) images. Though one can roughly regard each cell (row) as a sentence or an image patch, the 20,000 genes is a vast number compared to the typical sequence length, which is mostly a few hundred and no more than a few thousand (Qiu et al., 2020; Han et al., 2021). Thus directly applying existing transformer architecture will not work.

Second, scRNA-seq matrices are highly sparse. Although the number of genes is large, only a small fraction of the gene expression data is non-zero. The abundance level of RNA for each gene is measured by counting the unique molecular identifier (UMI) reads in scRNA-seq experiments (Jovic et al., 2022; Chen et al., 2019). However, many genes exhibit low UMI counts due to limited probing efficiency. In the data used for the pre-training task, the proportion of zero entries reaches 90%. Therefore, treating scRNA-seq data as an image and utilizing a convolutional neural network to extract features is not feasible, as it introduces huge number of redundant computations for sparse positions. A new architecture that can exploit the sparsity of scRNA-seq matrices is essential to allow for the efficient processing of these data.

Third, the gene expression values in scRNA-seq data are continuous scalars, which typically indicate similar gene activity when they have similar values. To transform these scalars into high-dimensional tokens in the data matrix, a representation that preserves the continuous semantics is needed. This requirement is different from NLP where tokens are discrete categorical values and CV where tokens are derived from patches. Manually discretizing the gene expression values is challenging as it is difficult to determine the best discretization thresholds and can result in sharp changes in category assignment near the thresholds. A learned discretization approach or learnable representation, such as the one proposed in (Gorishniy et al., 2022), is ideal for preserving the continuous semantics of the gene expression values.

Taking into the above three major features, we design a new architecture as described in the next section.

## 3 xTrimoGene Architecture

xTrimoGene is a highly efficient framework for pre-training large-scale single-cell RNA-seq data (illustrated in Figure 1). The training process is based on a regression masked task, aimed at accurately recovering masked values in the expression matrix. Notably, a specific optimized asymmetrical encoder-decoder framework is employed to accelerate the learning of sparse matrices. This is achieved by feeding only the unmasked non-zero positions (less than 10% of the full length) into the encoder, while the largely masked and zero positions are input into a lightweight decoder with a reduced number of layers and attention heads. This unique architecture significantly reduces computational costs and training time. In addition, a novel auto-discretization strategy is introduced to project continuous expression values into a latent embedding space. Instead of rounding to the nearest integer, values are directly mapped to the latent space allowing for the representation of closely-related values. The xTrimoGene framework consists of the following components:

1. **Masking**: A portion of the normalized gene expression matrix *V* input to the model is masked for prediction, including both zero and non-zero positions. c denotes cell sample size, and *n* denotes gene number (19,264 in our setting).
2. **Filtering**: The masked and zero-valued embeddings are filtered out, yielding a variable-length sequence of valuable information that is prepared for encoding.
3. **Padding**: The remaining unmasked positions are aligned with padding tokens, resulting in a much smaller unmasked-only matrix *V_unmasked_*. *m* denotes the maximum length of the unmasked sample.
4. **Embedding**: Expression value and gene embeddings are separately projected. *d* denotes the dimension of the embedding. The expression embedding is calculated through an auto-discretization mapping, which involves a sequential linear projection layer. The gene embedding is retrieved from a randomly initialized lookup table.
5. **Combining Expression and Gene Embeddings**: The expression and gene embeddings (*E* and *G*) are element-wise added to form the input embedding, which is then fed into the encoder of the model.
6. **Encoding**: The sum of the embeddings is input into the encoder, which implements self-attention mechanisms using a Transformer-like architecture.
7. **Extending masked and zero embeddings**: The intermediate encoder embedding *I_encoder_* is combined with embeddings for masked and zero-value positions.
8. **Decoding**: The combined embeddings are processed by the decoder, utilizing self-attention mechanisms instead of the typical casual attention used in NLP decoders.
9. **Loss Computation**: Decoder embedding is projected to model output with a MLP layer. The mean squared error (MSE) loss is computed between the predicted masked values from the model and their corresponding ground truth values.

**Figure 1:**
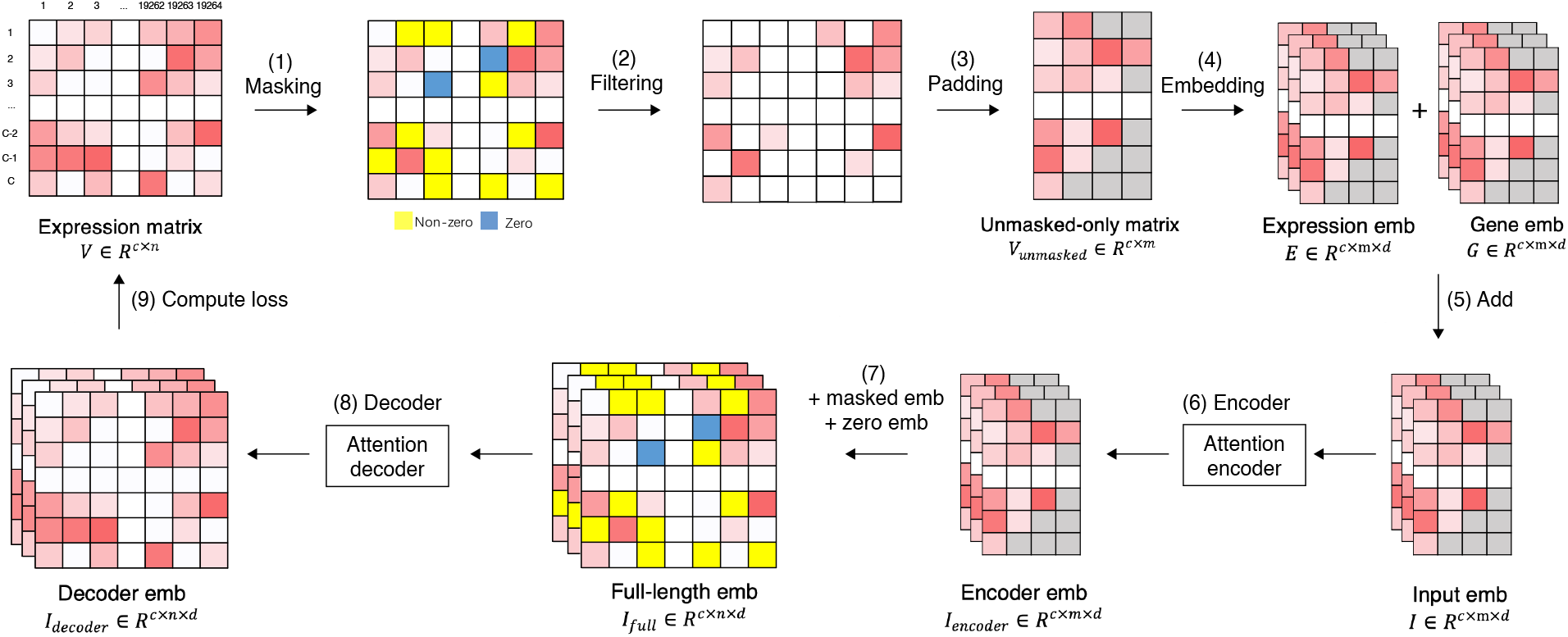
The xTrimoGene Framework: (1) Random positions (including both zero and non-zero values) are masked for prediction. (2) Masked and zero-valued positions are filtered out. (3) Remaining unmasked positions are aligned with padding tokens (grey) to ensure maximum length consistency within a batch. (4) Gene expression values and gene embeddings are separately projected into embeddings. (5) These two embeddings are element-wise added. (6) The resulting input is fed into the encoder. (7) The intermediate encoder embedding is combined with embeddings for masked positions and zero embeddings. (8) This combined representation is then fed into the decoder. (9) Decoder embedding is projected to model output with a MLP layer. The MSE loss is calculated between the model output and ground truth values for the masked positions.

In the subsequent sections, we will present the crucial components in detail.

### 3.1 Input expression matrix

The input to the model is a normalized cell by gene expression matrix 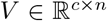. The normalization is performed using the standard pipeline from the Scanpy (Wolf et al., 2018). This study focuses on human scRNA-seq data and considers only protein-coding genes, resulting in a substantial expression matrix with a sample size of close to 5 million cells and 19,264 genes. Further information on data collection and processing can be found in Appendix 7.1.

### 3.2 Details of encoder

The scRNA-seq data is characterized by its high sparsity, with cell information largely concentrated in the non-zero expression values. Thus, the encoder is designed to focus only on the non-zero part of the unmasked matrix, *V_unmasked_*. The encoder is based on a traditional multi-head attention transformer and takes the combination of value embedding, E, and gene embedding, *G*, as its input, 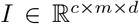. The value and gene embeddings are similar to the word and positional embeddings in natural language modeling, respectively. The value embedding, E, is generated using the auto-discretization strategy discussed previously, while the gene embedding, G, is retrieved from the embedded vocabulary.

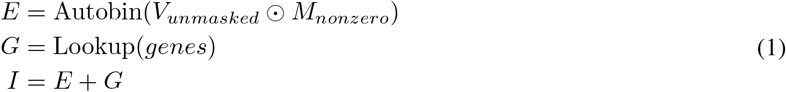

Then the encoder processes the input embeddings *I* and generates the high-level gene representations 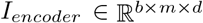 via the multi-head attention mechanism.

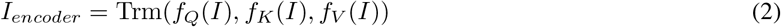

where *f_Q_*, *f_K_*, *f_V_* are the project functions. *Trm* denotes the Transformer block.

It is worth emphasizing that our encoder only operates on a subset of genes, reducing the length of the processed sequence to 1/10 of the original. This allows the full-length transformer to be used without any computational approximations.

### 3.3 Details of decoder

Unlike the encoder which focuses on the main information (non-zero expression values) in the cells, the decoder in the system performs full-length feature abstraction and extraction. The input to the decoder, *I_full_*, comprises three token types: the output from the encoder, *I_encoder_*, the genes with zero expression embs *I_zero_*, and the mask token embs *I_masked_*. Out of these tokens, genes with zero expression make up 90% of all tokens. The gene embeddings are concatenated with all of these tokens to provide the decoder with gene-specific information for the corresponding mask tokens, then followed by a Full connection layer.

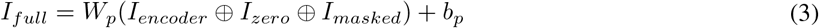

where ⊕ represents the concatenation operation, and *W_p_* and *b_p_* are learnable parameters that project the decoder’s embedding size.

The decoder in the framework is optimized for long-sequence attention calculations and employs the Performer architecture as its backbone. The decoder transforms the input *I_full_* into final gene-level embeddings, 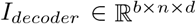, and predicts the masked values through a shared linear layer, 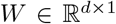, applied across all genes. The operations are expressed as follows:

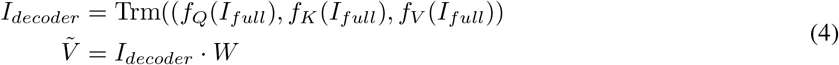

The decoder has a smaller model size compared to the encoder, with a smaller embedding size, fewer attention layers, and fewer attention heads. For instance, in the largest model configuration, the layer depth ratio between the encoder and decoder is 2:1 (12:6) and the head number ratio is 1.5:1 (12:8). Similarly, the principle of asymmetric encoder-decoder design has been proven powerful in masked autoencoders (MAE) (He et al., 2021), which is tailored for CV data pre-training. Unlike MAE, xTrimoGene utilizes the biased masking strategy to avoid the learning process being dominated by zero tokens. Though the scRNA-seq data is distinct from images, our results show that the performance gains of xTrimoGene are comparable to those of MAE, with more efficient training and better downstream task performance.

### 3.4 Auto-discretization strategy

The expression value 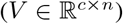 is transformed into a hidden embedding (*E*) for addtion with the gene embedding (*G*) via an auto-discretization block. This block uses a random look-up table 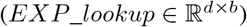, where *d* is the embedding dimension and *b* is the number of tokens (default 100). The expression value is first transformed with a linear layer (*v*_1_ = V · *w*_1_), where *w*_1_ is the weight vector, then passed through a leaky ReLU activation (*v*_2_ = *leaky_relu*(*v*_1_)). A cross-layer projection (*v*_3_ = *w*_2_ · *v*_2_ + *α* · *v*_2_), with weight vector *w*_2_ and scaling mixture factor *α*, follows. The bin weights are normalized using the Softmax function (*v*_4_ = *softmax*(*v*_3_)). The final output is a weighted combination of individual embeddings from the look-up table (*output* = *EXP_lookup* · *v*_4_), where the weights are learnable parameters.

To validate the effectiveness of the expression value projection, we conducted an analysis of viewing the weight distribution pattern for continuous values. Our results showed that the normalized weight distribution of the close values exhibited smooth transitions and that of the distant values being clearly distinguishable (Figure 2). This supports the conclusion that the auto-discretization strategy effectively represents continuous values with high resolution while preserving relatively rich meaning.

**Figure 2:**
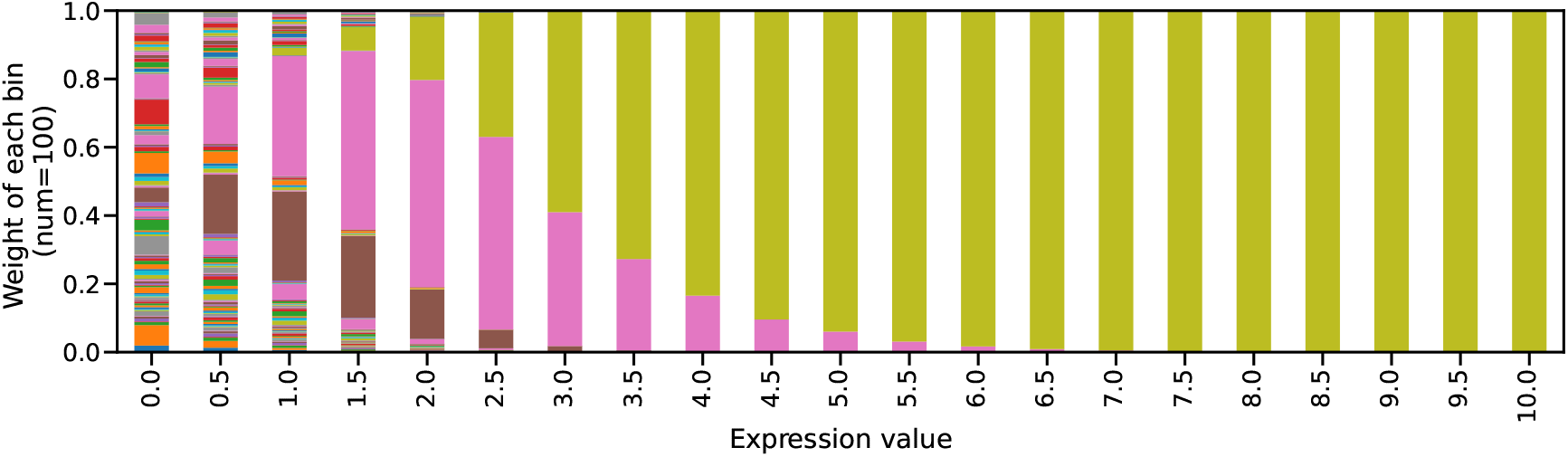
Weight distribution across bins for various expression values. The auto-discretization strategy was applied to each expression value in the range from 0 to 10, producing a corresponding weight vector with a length equal to the number of bins (100 in this case). The weight vectors were normalized to sum to 1 and visualized as stacked plots.

We also compared the performance of the proposed auto-discretization strategy with three other discretization methods: (1) Round bin with zero, in which values are rounded to the nearest integer, and zeros are kept as it is, (2) Up bin without zero. Values greater than zero are converted to the nearest ceiling integer, while zero is represented as individual 0. (3) Equal bin. All the values fall into a fixed percentage interval, which is calculated by value distribution and frequency. We evaluated the different strategies on a standard cell clustering task (refer to Appendix 7.2 for details) and found that the proposed auto-discretization strategy outperformed the others (as shown in Figure3), demonstrating the importance of high-resolution projections in handling expression values.

**Figure 3:**
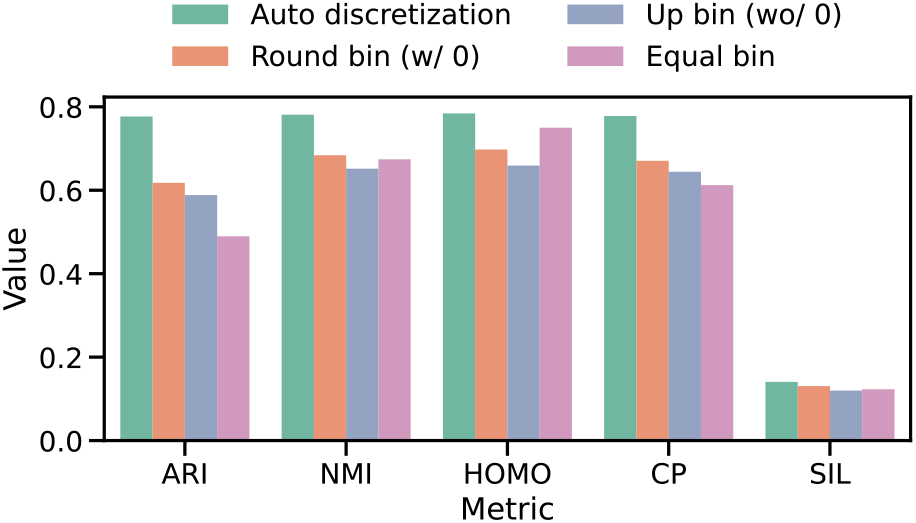
Perofrmance comparison between auto discretization strategy and other binning methods for expression value projection. The cell clustering task is evaluated and five metrics are displayed. ARI for Adjusted Rand index, NMI for Normalized Mutual Information, HOMO for Homogeneity, CP for Completeness and SIL for Silhouette Coefficient. The details of the definition and calculation for all metrics are referred to in Appendix 7.2.

## 4 Training Strategy

We now explain the strategy used to train the asymmetric encoder-decoder transformer.

### 4.1 Regression masked task

The traditional masked language task is a multi-class classification problem, where the predicting target is a single token with limited, naturally distinct categories. In contrast, the normalized gene expression value is a continuous scalar. To fit the data property, we modify the pre-trained learning objective to a regression task, aimed at recovering the absolute value of the masked positions. The loss function employed is the Mean Square Error (MSE) between the ground truth and the predicted values:

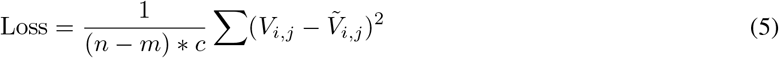

where *n* represents the number of all genes, *m* represents the maximum length of the unmasked positions in a sample, and c represents the number of cells. To evaluate the efficacy of this modification, we compared the regression setting with the classification setting on the cell clustering task. The results indicate that the regression model outperforms the classification model (Figure 4), providing evidence of the benefits of learning a more fitted representation.

**Figure 4:**
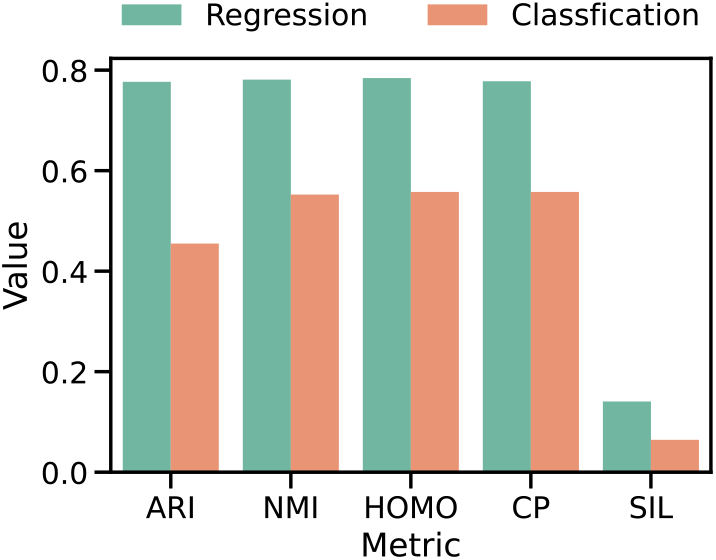
Performance of pre-trained models with different task mode, including regression and classification setting. The cell clustering task is evaluated. Detailed descriptions can be referred to the main text.

### 4.2 Masking strategy

We mask both non-zeros and zeros positions though the scRNA-seq expression matrix is highly sparse (where zero percentage is usually over 90%). Apart from limited experimental protocol resolution, some zeros represent the true extremely low expression level. This type of zeros is informative to illustrate how the gene abundant behaves inside the cell. Learning zero embedding is also helpful in building a complete gene-gene interaction map. As the zero positions percentage is much higher than non-zero positions, the masked ratio can’t be the same for the two types. Otherwise, the model tends to predict all zeros and still obtains a low error level. We propose to mask an almost equal number of positions for zero and non-zeros positions, as illustrated in Table 1. The setting enforces the model to learn embeddings for all values and not to be dominated by zero representation.

**Table 1:** Masking strategy for gene expression matrix. The gene expression matrix is masked by selecting a predetermined number of positions for prediction. ~ 1,100 positions, including ~600 non-zero and ~540 zero expressions, are masked in a matrix with ~20,000 genes. The performance of the model is evaluated using Mean Squared Error (MSE) loss on these masked positions.

| Value | Masked | Unmasked | Total |
| --- | --- | --- | --- |
| $\neq 0$ | 600 | 1,400 | 2,000 |
| $= 0$ | 540 | 17,460 | 18,000 |
| sum | 1,140<br>(5.7%) | 18,860<br>(94.3%) | 20,000<br>(100%) |

The recovery of masked tokens in NLP is challenging due to the fact that word comprehension relies heavily on long-range interactions rather than local context. Accurate inference of the missing tokens can be achieved at low masking ratios (15%) where the information in the entire sentence is still relatively redundant and encoded by the unmasked tokens. We investigated the density of information needed for the scRNA-seq regression task by training models with different masking ratios (for non-zero values, the ratio was set 10 times higher than for zero values) ranging from 15% to 90% with a 15% interval. The models were then evaluated on the cell clustering task, with the results showing that performance improved first and then degraded as the masking ratio increased. When the masking ratio was close to 30%, the majority of metrics reached a peak (Figure 5). These results suggest that the scRNA-seq expression vector contains more redundant information than a sentence and highlight the role of hidden regulations between genes in constraining the inference of expression values.

**Figure 5:**
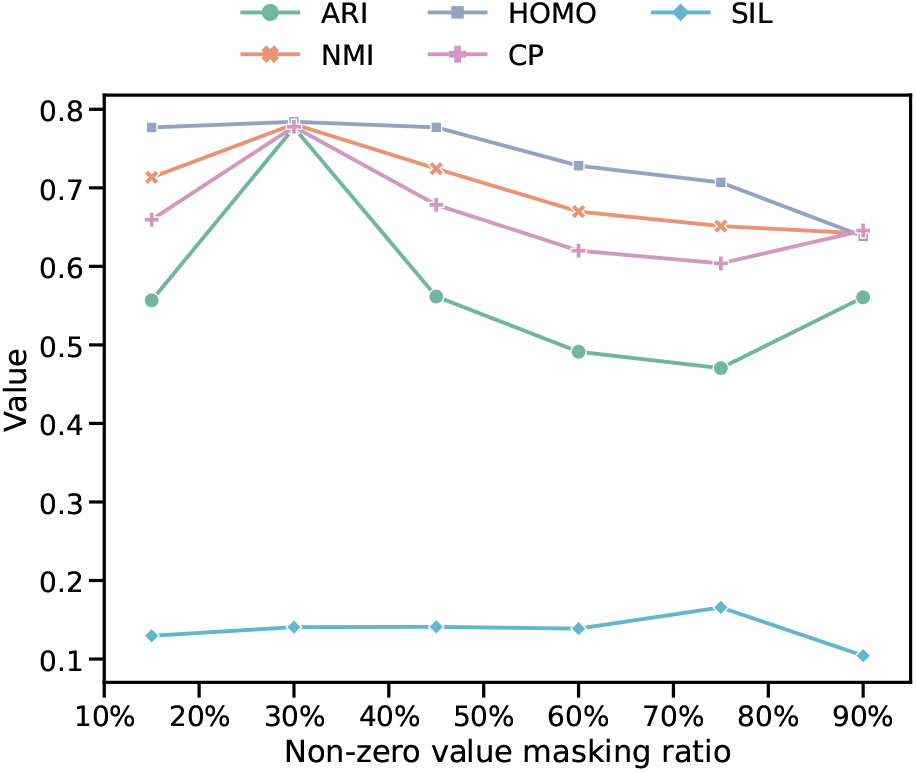
Model performance under different mask ratios of non-zero values. The cell clustering task is evaluated.

### 4.3 Acceleration strategy

The attention mechanism in masked language modeling is computationally expensive for long sequences, as time and space complexity grows quadratically along with sequence length. Though multiple attention architectures have been proposed to reduced the complexity to near linear, it’s still slow to train large models with billions of data. We have adopted multiple techniques to boost the training speed as following.

Since FP16 or BFLOAT16 Tensor Core have twice the computational throughput compared to TF32 on NVIDIA Ampere GPU, and, additionally, FP16 training also reduces residual memory consumption, xTrimoGene was conducted mainly with mixed-precision training strategy to optimize computational efficiency.

Distributed Data Parallelism is another training strategy used in our work, which handles large corpus on HPC clusters. In our setting, one single Ampere GPU provides sufficient amounts of memory for one model replica of billions of parameters performing forward and backward passes, and gradient accumulation is used to raise effective batch size to enhance large model training.

To scale up the model size, ZeRO-DP stage two (Rajbhandari et al., 2020) and checkpointing (Chen et al., 2016) techniques are experimentally tested in our setting. The results verified that both strategies reduce the model and residual state memory without expanding training time too much.

## 5 Experiments

Next we will explain our experimental settings and results.

### 5.1 Dataset description

All the scRNA-seq data are collected from Gene Expression Omnibus (GEO) repository with a keyword searching and data retrieval process. Then the downloaded count matrices are processed with a unified pipeline, including reference gene list mapping, quality control and normalization (see Appendix 7.1 for details). In total, we curated about 5 million scRNA-seq data for training. The full data set is randomly split into train, validation and test sets with ratio of 96:2:2.

### 5.2 Computational efficiency

We quantitatively compared the training cost of xTrimoGene with other two decoder-only models, including full-length attention Transformer and kernel-based approximation Performer (scBERT). For an apple-to-apple comparison, three models are set to approximately 10 million trainable parameters and trained on 5 million samples over 5 epochs. We calculated the corresponding FLOPs, where matrix multiplication operations are considered only. We observed that total FLOPs for Performer (scBERT) decreased to 10% of native Transformer (see Table 2). Notably, xTrimoGenes runs a 3-times faster than Performer. The results validate the efficiency of xTrimoGene, which is readily adapted for large-scale data pre-training.

**Table 2:**
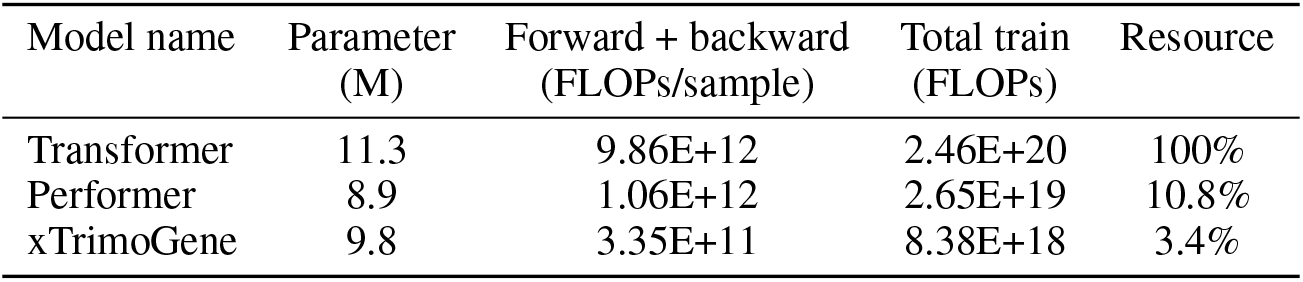
Computational efficiency comparison between different algorithms. Transformer and Performer are evaluated with the decoder-only framework. Performer is the default attention block in scBERT. All models are set to train on a 5 million data set and for 5 epochs. The resource column is normalized by the Transformer row.

| Model name | Parameter (M) | Forward + backward (FLOPs/sample) | Total train (FLOPs) | Resource |
| --- | --- | --- | --- | --- |
| Transformer | 11.3 | 9.86E+12 | 2.46E+20 | 100% |
| Performer | 8.9 | 1.06E+12 | 2.65E+19 | 10.8% |
| xTrimoGene | 9.8 | 3.35E+11 | 8.38E+18 | 3.4% |

### 5.3 Scalability

The Deep Learning community has shown significant interest in the scalability of proposed models (Kaplan et al., 2020; Brown et al., 2020). Vanilla Transformer models are challenging to scale due to their computational time and resource requirements, which increase quadratically with model size. Varieties of attention mechanisms have been proposed to accelerate training speed, a critical factor for model scaling. However, increasing model size is not always consistent with performance gains. For example, (Tay et al., 2022) showed that the Performer model struggles with scale-up. Although the Performer-large model achieved slightly lower perplexity than the Performer-base model during upstream pre-training, it obtains nearly no improvement on downstream tasks.

To test the scale-up ability of xTrimoGene, we further pre-trained three models across multiple compute regions and scales (e.g., from 3M to 100M parameters). The detailed hyperparameter setting is displayed in Table 3. The training curve clearly shows all models are steadily down to a lower loss when training steps increase (Figure 6). More importantly, xTrimoGene-100M model obtains a significant improvement over xTrimoGene-10M model, which is also superior over xTrimoGene-3M model. The results suggest xTrimoGene framework is robust to scale-up, making it possible and convenient to pre-train larger models with more data.

**Table 3:** Size and hyper-parameters of the pre-trained models. All models are set to train on a 5 million data sets and for 5 epochs.

| Model name | Parameter<br>(M) | depth | Encoder<br>heads | dim | depth | Decoder<br>heads | dim |
| --- | --- | --- | --- | --- | --- | --- | --- |
| xTrimoGene-3M | 3 | 4 | 2 | 128 | 2 | 2 | 128 |
| xTrimoGene-10M | 10 | 4 | 8 | 256 | 2 | 4 | 256 |
| xTrimoGene-100M | 100 | 12 | 12 | 768 | 6 | 8 | 512 |

**Figure 6:**
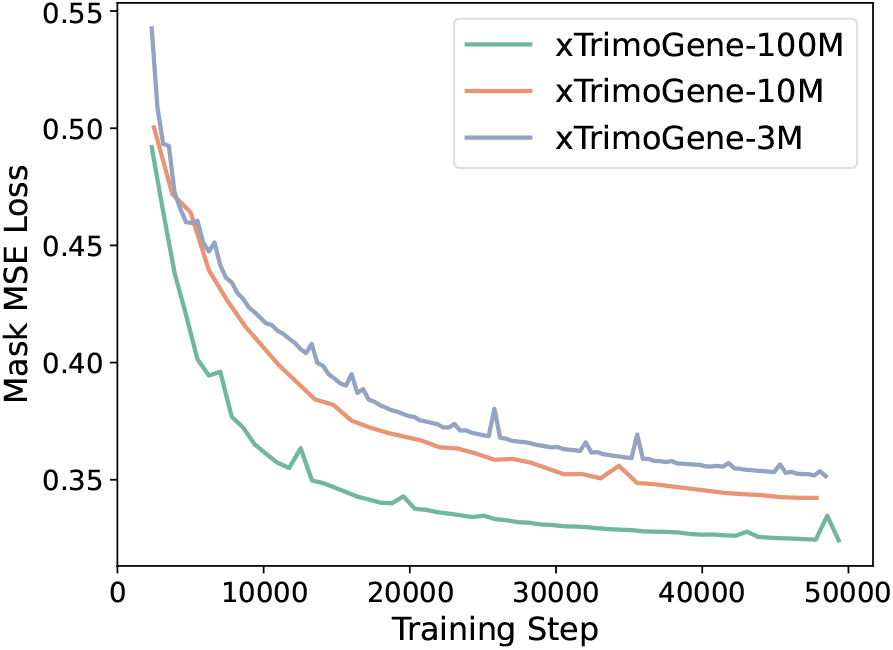
The learning curve of pre-trained xTrimoGene models with different parameter scale. The loss curve measures MSE for masked positions during the pre-training stage, and only the validation set is displayed.

### 5.4 Robustness on high sparse data

scRNA-seq data often exhibit varying levels of sparsity, thus it’s necessary to assess whether xTrimoGene is robust in handling different sparse data. To verify the robustness, we divided the test samples into subgroups based on cell type and calculated the sparsity level (i.e., percentage of zero values in the expression matrix) and Pearson correlation coefficient between the predicted and actual values. Our results reveal that the correlation gradually decreases as the sparsity level increases, as expected (Figure 7A). However, the correlation remains above 0.8 even when the sparsity level reaches 96% (Figure 7A), indicating the robustness of xTrimoGene. We also compared xTrimoGene’s performance with Performer, and found that xTrimoGene consistently achieves a higher correlation across most subgroups (Figure 7B). These findings demonstrate that xTrimoGene is robust in handling highly sparse data and outperforms decoder-only architectures.

**Figure 7:**
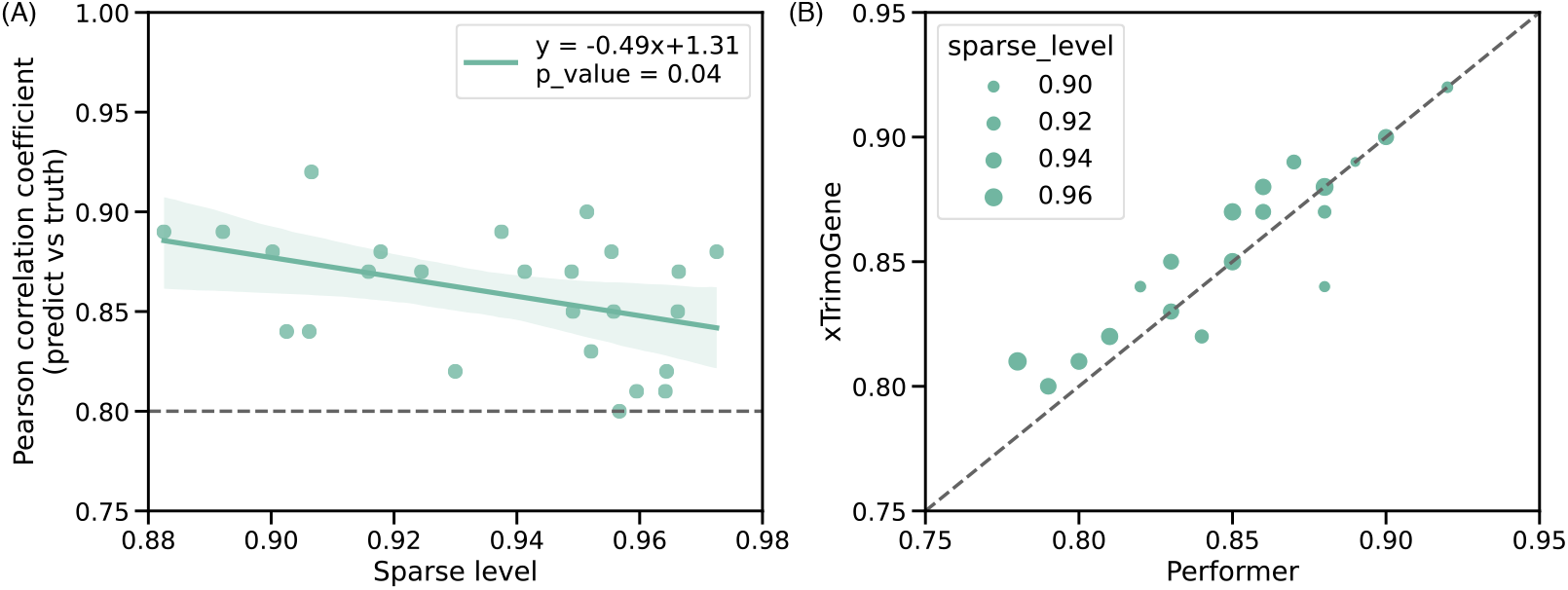
Comparison of performance for different sparse level data. (A) xTrimoGene performance for recovering masked values at different sparse levels. Each dot represents a subset defined by cell type. Sparse level is calculated as the ratio between zero value percentage. Pearson correlation coefficient metric is calculated on masked positions. (B) Performance comparison of xTrimoGene and Performer while recovering masked values at different sparse levels. Dot has the same meaning as (A) but the dot size is proportional to the sparse level. Both the x and y axis denotes the Pearson correlation coefficient metric for a particular algorithm.

The performance of the encoder-decoder and decoder-only architectures have been comparatively analyzed in the NLP domain, with the former demonstrating effectiveness in language comprehension and the latter in context generation. Apart from comparison on masked value recovery, we further evaluated xTrimoGene against decoder-only Performer on the cell clustering task. The results demonstrate that xTrimoGene achieves superior performance, reaffirming its proficiency in latent embedding extraction (Figure 8).

**Figure 8:**
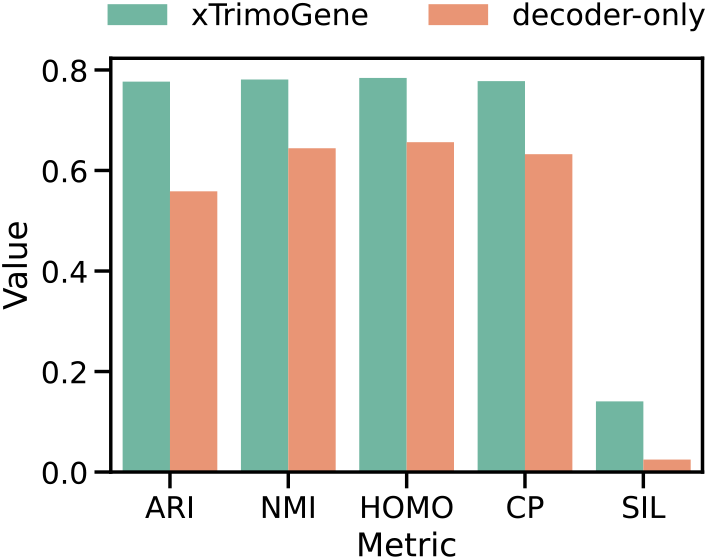
Comparison of performance for xTrimoGene framework and decoder-only framework. Cell clustering task is evaluated.

### 5.5 Explainable case study of biological representation

The pre-training task endows the model with the ability to capture the relationship between genes. This can be demonstrated by feeding the model with a biologically meaningful binarized vector. Specifically, we collected the cell type-specific marker genes of B cells and Fibroblast from PanglaoDB (Franzén et al., 2019), and selected the top 50 genes with high ubiquitousness index, respectively. We initialize two zero vectors with lengths equal to total gene number. Then the value for corresponding cell maker genes are set to 1. Next, the two simulated binary expression vectors are input to the xTrimoGene. The top 500 highly expressed genes among the model outputs are selected as activated genes. For the two gene sets, we found 427 genes are common and 73 genes are cell-type specific. We conducted KEGG pathway enrichment analysis on these 73 genes via the enrichR online tools (Chen et al., 2013; Kuleshov et al., 2016). As shown in Figure 9, the highly expressed genes corresponding to B cell vectors are enriched in JAK-STAT, PD-1 and other human immunity and cancer-related pathways (Papin and Palsson, 2004; Han et al., 2020), while the highly expressed genes corresponding to Fibroblast are enriched in cardiac fibrosis, liver fibrosis and other fibrosis diseases related pathways(Deb and Ubil, 2014; Wong and Adams, 2018). This illustrates the ability of our model to infer the hidden relation between genes in different cell types from very limited gene expression information.

**Figure 9:**
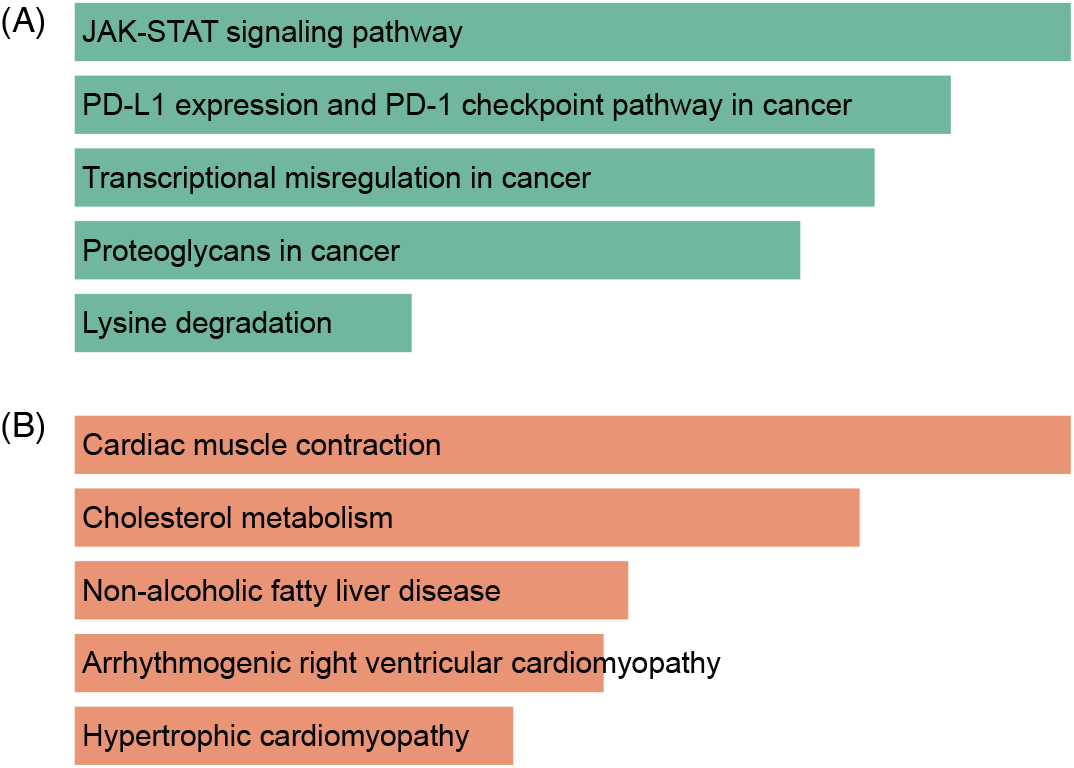
KEGG enrichment analysis of top expressed genes activated by particular marker genes. Bar plot (A) and (B) shows the enriched pathway term on B cell and Fibroblast marker gene, respectively.

### 5.6 Evaluation on downstream tasks

Analyzing scRNA-seq data has greatly advanced our understanding on cell biology, which process is often accelerated by growing algorithms. Currently, multiple tasks have been established to evaluate different models, including well-defined cell type annotation and recently developed perturbation response prediction tasks. We first assessed the performance of xTrimoGene on these single-cell tasks. Additionally, we explored the potential application on bulk RNA-sequencing data, with a focus on synergistic drug combination prediction.

#### 5.6.1 Cell type annotation

First, we evaluated xTrimoGene’s performance on cell type annotation task with Zheng68K (Zheng et al., 2017) dataset, which has been widely benchmarked. The dataset is a mixture of 12 unbalanced cell types, making it difficult to distinguish and appropriate for model assessment. We compared the xTrimoGene against other several methods, including scBERT, ACTINN (Ma and Pellegrini, 2020) and Scanpy (Wolf et al., 2018). For xTrimoGene model, we added a maxpooling layer and a linear layer to predict cell type labels with fine-tuning mode. For other methods, we followed their instruction with the default parameter setting. More details can be found in the Appendix 7.3. We observed that xTrimoGene achieves a high Precision (0.7335) and F1 score (0.7354), surpassing all the other methods (Table 4). The results indicated xTrimoGene learns a well-represented cellular embedding by simply aggregating contextual gene embedding.

**Table 4:**
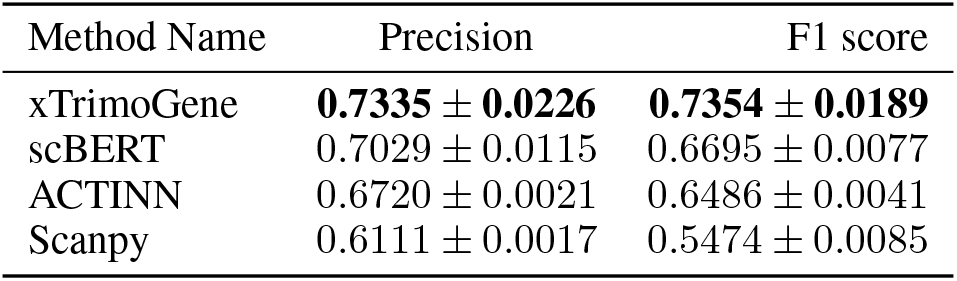
The cell annotation performance on the Zheng68K dataset. xTrimoGene is evaluated with 10M parameter model.

| Method Name | Precision | F1 score |
| --- | --- | --- |
| xTrimoGene | **0.7335 ± 0.0226** | **0.7354 ± 0.0189** |
| scBERT | 0.7029 ± 0.0115 | 0.6695 ± 0.0077 |
| ACTINN | 0.6720 ± 0.0021 | 0.6486 ± 0.0041 |
| Scanpy | 0.6111 ± 0.0017 | 0.5474 ± 0.0085 |

#### 5.6.2 Perturbation response prediction

Recently, perturb-seq technology was established to screen gene expression response given pooled perturbations at single-cell level (Dixit et al., 2016). Comparing before and post-perturbation data, downstream differential expression (DE) analysis leads to the identification of genes critical to disease progression. Several algorithms have also been developed to systematically predict perturbation effects (Roohani et al., 2022; Lotfollahi et al., 2021) at single cell level, i.e., what is the expression value of genes after perturbation. Typically, the gene expression profile before perturbation and perturbed target information are separately encoded and subsequently combined for joint learning.

As xTrimoGene is capable of generating representative embedding for expression profiles, we tested whether xTrimoGene is beneficial for the task. We compared the native GEARS model with and without incorporating embeddings from xTrimoGene. While utilizing xTrimoGene, the normal state (before perturbation) gene expression profile is fed into xTrimoGene and we obtained the context embedding, which replaces raw expression value input in GEARS model (refer to Appendix 7.4 for detailed description). All the other settings remain unchanged. The evaluated data set (Norman et al. (Norman et al., 2019)) contains both single and double gene perturbation, we thus assess the performance across different perturbation levels. As shown in Figure 10, GEARS with xTrimoGene embedding scores a lower MSE (decreased 14.8%) for top20 differential expressed genes across all perturbation scenarios. Notably, the tendency is consistent across different perturbation levels, regardless the perturbed target is seen or not. The results demonstrated that the pre-training strategy empowers xTrimoGene to capture constraints under various circumstances, including post-perturbations. The application further proved the efficacy and potential of xTrimoGene to boost scRNA-seq based tasks.

**Figure 10:**
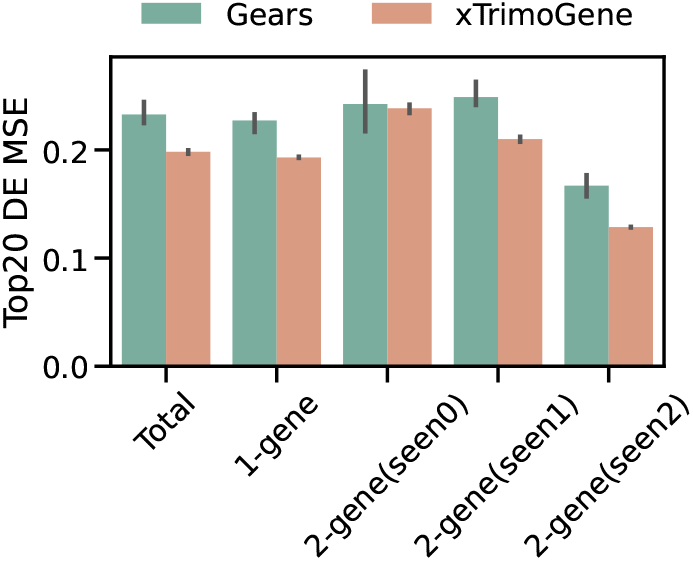
The MSE of the top 20 deferentially expressed (DE) genes given by different models on perturbation response prediction. The top20 DE genes are calculated between the before and post-perturbation expression profile. “Total” denotes evaluating all test perturbation set. “1-gene” denotes evaluation on the single gene perturbation subset, where the perturbed target is not seen in the training set. “2-gene” represents the sub-test set for perturbing two genes simultaneously. “seen0”,”seen1” and “seen2” denotes zero, one or two perturbed targets are not seen in the training set, respectively. Black line denotes 95% confidence interval.

#### 5.6.3 Synergistic drug combinations prediction

Drug combination therapy has been proven effective in clinical experiments (Mokhtari et al., 2017). The therapy evaluates how patients or cells respond to a drug combination intervention. Recently, high-throughput screening technology has been developed to measure the drug synergy effect at a large scale, which is promising in prioritizing drug candidate. However, the generated wet-lab experimental data only covers a tiny search space of possible drug combinations. Multiple models have been proposed to accelerate predicting the synergistic landscape of drugs (Preuer et al., 2017; Wang et al., 2022). For instance, DeepDDS integrates genomic expression profile and drug chemical information with graph-based neural network, greatly improving the prediction performance.

The bulk RNA expression profile represents the normal state of a population of cells, which implicates the regulations between genes. The expression profile is always encoded with different blocks (e.g., sequential linear layers) to extract the high-level representation. We further explored whether xTrimoGene is able to generate good latent embedding for bulk expression data. Similar to the perturbation prediction test, we adapted xTrimoGene to DeepDDS with the intermediate context embedding (refer to Appendix 7.5 for detailed description). We also included DeepSynergy and Random Forest (RF) for comparison. As illustrated in Figure 11, utilizing embedding from xTrimoGene model outperforms all the other models. The result proved xTrimoGene can accurately capture cell level representation, even for bulk sequencing data. This also opens the avenue for xTrimoGene to be applied across other biological modeling tasks, especially where bulk level transcriptome data is available.

**Figure 11:**
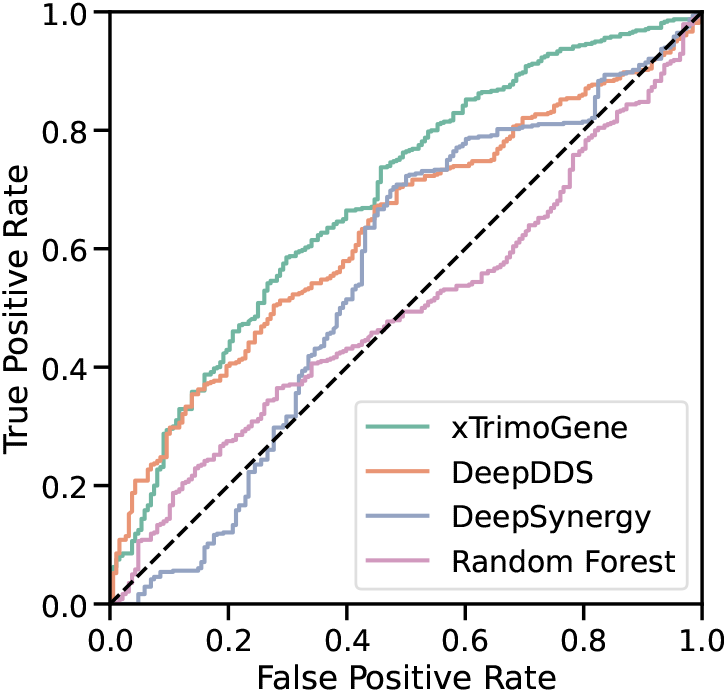
ROC curve of different models on drug combination synergy prediction task. xTrimoGene denotes replacing the raw expression profile with context embeddings in DeepDDS framework and others remain unchanged. Refer to Appendix 7.5 for more details.

## 6 Conclusion

xTrimoGene is a new, efficient framework for learning scRNA-seq data. It proposes an asymmetric encoder-decoder framework that takes advantage of the sparse gene expression matrix, and establishes the projection strategy of continuous values with a higher resolution. The results show that xTrimoGene is scalable and performs well on tasks like cell type annotation, perturbation response prediction, and synergistic drug combination prediction. The experiments demonstrate the efficacy of pre-training in single-cell biology. xTrimoGene has been integrated into BioMap’s singlecell analysis platform, functioning as a fundamental and essential model (as depicted in the supplementary material’s Figure 12). The codebase, pre-trained model, and accompanying training and evaluation pipelines will be publicly available for access on GitHub ^1^ soon. In the future, with the increase of data, larger pre-trained models are expected to drive more advancements in various downstream task learning.

**Figure 12:**
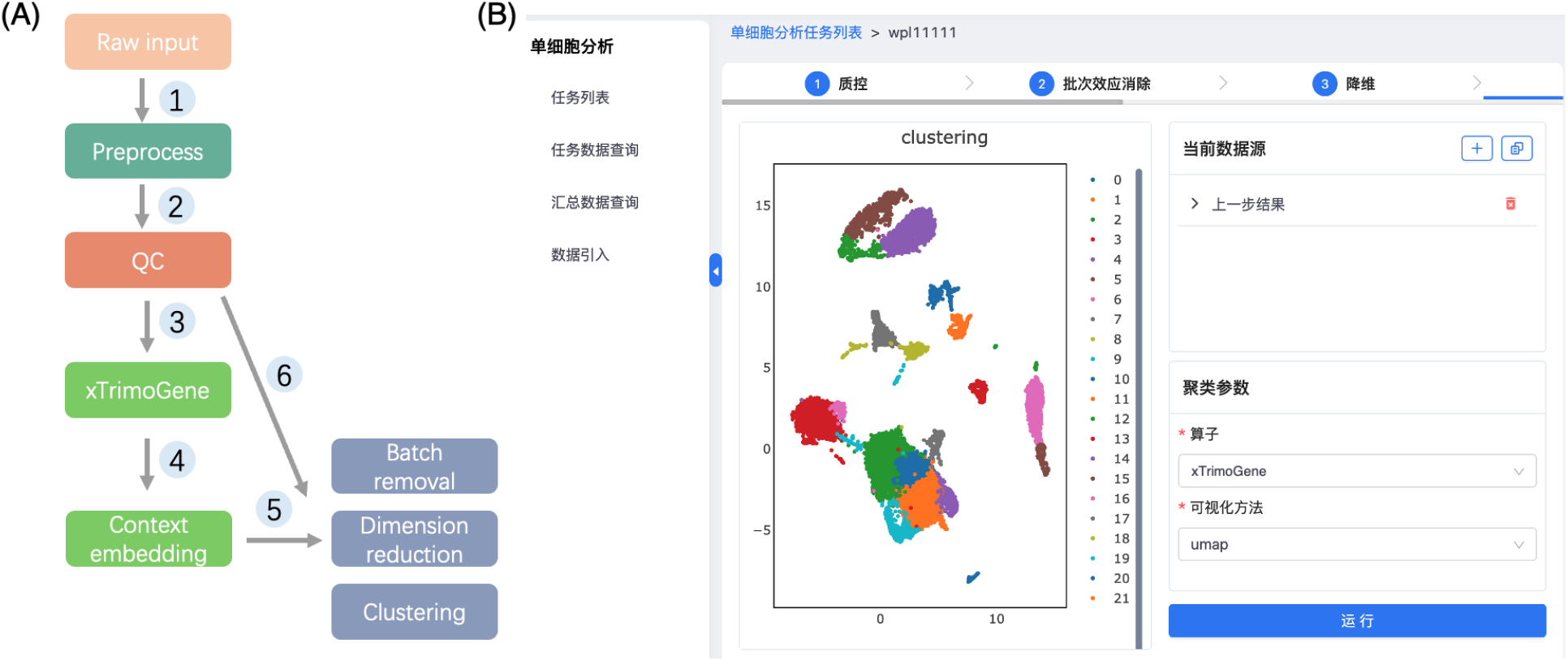
The deployment of xTrimoGene model on a website is depicted in this figure. Figure A shows the overall pipeline, which includes the following steps: (1) User-uploaded raw input undergoes preprocessing and filtration through (2) quality control, (3) feeding the processed data into xTrimoGene for (4) context embedding extraction. The model supports multiple downstream applications such as (5) cell clustering, dimension reduction, and batch removal. The extracted expression profile can also be directly utilized by other algorithms. Figure B provides a snapshot of a clustering task in action using xTrimoGene’s context embeddings.

## 7 APPENDICES

### 7.1 scRNA-seq data collection and processing

Recently, scRNA-seq data is rapidly accumulated and mostly has been uploaded to Gene Expression Omnibus (GEO) repository (https://www.ncbi.nlm.nih.gov/geo/). We major collected data from GEO and processed data with a unified pipeline.

#### Downloading data and preparing raw count matrix

We first search scRNA-seq related data sets in GEO with multiple keywords, including “scRNA-seq”, “single cell RNA-seq”, “single cell RNA-seq sequencing”. The search processes return a list of GSE ID from different studies. After removing duplicated GSE ID, we downloaded the particular expression or count matrix. Most of the samples provide a raw count matrix. For samples with normalized expression matrices, we converted the matrix back to a count matrix. The conversion strategy is as follows: the minimal non-zero value in the whole normalized matrix is thought to have raw count 1, then all the other normalized values can be converted by scaling to this minimum value.

#### Matrix mapping to the reference gene list

After preparing all the count matrices, we mapped the matrix to our reference gene list. We downloaded the human protein-coding gene list (about 19,000) from the HUGO Gene Nomenclature Committee (HGNC, https://www.genenames.org/download/archive/), plus common mitochondrial genes, jointly constitute our full reference list (*n* = 19,264). For each count matrix, values of those genes not mapped in the reference list are filled with zero.

#### Quality control

To filter low-quality samples, we only keep samples with over 200 genes expressed (i.e., expression vector with non-zero value count > 200) for subsequent training and analysis.

#### Normalization

We followed the standard process in Scanpy (https://scanpy-tutorials.readthedocs.io/en/latest/pbmc3k.html) (Wolf et al., 2018) to obtain the normalized expression value. There are two steps: (1) for each sample normalize the library size to 10,000. (2) scale the values into a log space.

### 7.2 Clustering task evaluation and data sets

Cell clustering is an essential task for single-cell research, reflecting the ability of cell embeddings to remove noise and preserve biological signals. In the ablation experiments, we benchmarked the models’ clustering performance on two cell-type annotated datasets.

#### PBMC

This dataset is processed by Scanpy python library (Wolf et al., 2018) and contains 2,638 cells. The cell types are annotated by the human with known markers and cover the major immune cells including B cells, CD4 &CD8 T cells, Monocytes, Dendritic cells, Megakaryocytes and NK cells.

#### Experiments and Evaluation Metric

For every single cell, the expression values are fed into the model and the max pooling layer is applied across all genes’ output embedding to get a cell embedding. We then perform the usual single-cell clustering analysis step on these embeddings: 1) build the neighboring graph based on these embeddings 2) use the Leiden algorithm to cluster the cell into groups. Since the Leiden algorithm requires resolution rather than the number of clusters, we used a dichotomy method to find an optimal resolution reaching the number of cell types given in the dataset.

Several evaluation metrics are applied to access the performance of the clustering results in different aspects, including: Adjusted Rand index (ARI), Normalized Mutual Information (NMI), Homogeneity(HOMO), Completeness (CP) and Silhouette Coefficient (SIL). All these metrics are the higher the better. ARI and NMI measures the similarity of the clustering results from the statistics and information entropy theory view, respectively. HOMO and CP are intuitive metrics using conditional entropy analysis. HOMO measures how much the sample in a cluster are similar, and CP measures how much similar samples are put together. SIL measures the similarity of the embeddings to its cluster member compared to other clusters.

### 7.3 Cell type annotation task

We downloaded the Zheng68K expression matrix dataset from (Zheng et al., 2017) and mapped the matrix to our reference gene list. Then, the dataset is split into training, validation and test sets with a ratio of 8:1:1. All the methods are trained on the training set and the best model is selected according to the performance on validation set. Evaluation metrics (macro F1-score and marco precision) are calculated for individual testing set.

In training process, the expression matrix is fed into the encoder of the xTrimoGene model and the gene embedding is obtained. Then we used a max-pooling layer to aggregate all gene embeddings into one cell embedding, and use a single linear layer to predict cell types from the embeddings.

### 7.4 Perturbation effect prediction

The Norman dataset is downloaded from a previous study (Roohani et al., 2022). The expression matrix data is mapped to our reference gene list. We reproduced results of GEARS with original codes and settings (https://github.com/snap-stanford/GEARS). All the data processing are the same as GEARS, including data split, pre-post sample pairing strategy and evaluation metrics calculation.

While training GEARS with xTrimoGene, the expression matrix is fed into xTrimoGene and intermediate context embedding is obtained. The context embedding is then input to the co-expression graph network branch, all the other parts remain unchanged.

### 7.5 Drug combination prediction

To test how xTrimoGene adapted to DeepDDS (Wang et al., 2022) for synergistic drug combination prediction, we first reproduced DeepDDS algorithm. Both data and original codes are downloaded from Github repository (https://github.com/Sinwang404/DeepDDs/tree/master). We use data in “new_labels_0.csv” file for training and “independent_set” for testing. The genomic expression data are all mapped to our reference gene list. Models are trained 5 times and evaluated on the testing set. For all metrics, the averaged value and the standard deviation are reported. We keep the overall framework of DeepDDS while testing xTrimoGene. The genomic expression matrix is fed to xTrimoGene and the intermediate context embedding is obtained. The embedding replaces raw expression profile for MLP branch input.

### 7.6 Website deployment of xTrimoGene model

xTrimoGene has been proven advantageous in gene representation and cell context embedding extraction. To facilitate its wide application for single-cell RNA-seq data analysis, we deployed the xTrimoGene model within Biomap corporation. On the website, the xTrimoGene is implemented as a standard operator and serves multiple downstream tasks, including cell clustering, dimension reduction and batch removal Figure (12). The interactive page is user-friendly and feasible to evaluate performance with rich visualizations.

